## supplementary material for "Structure of the central *Staphylococcus aureus* AAA+ protease MecA/ClpC/ClpP"

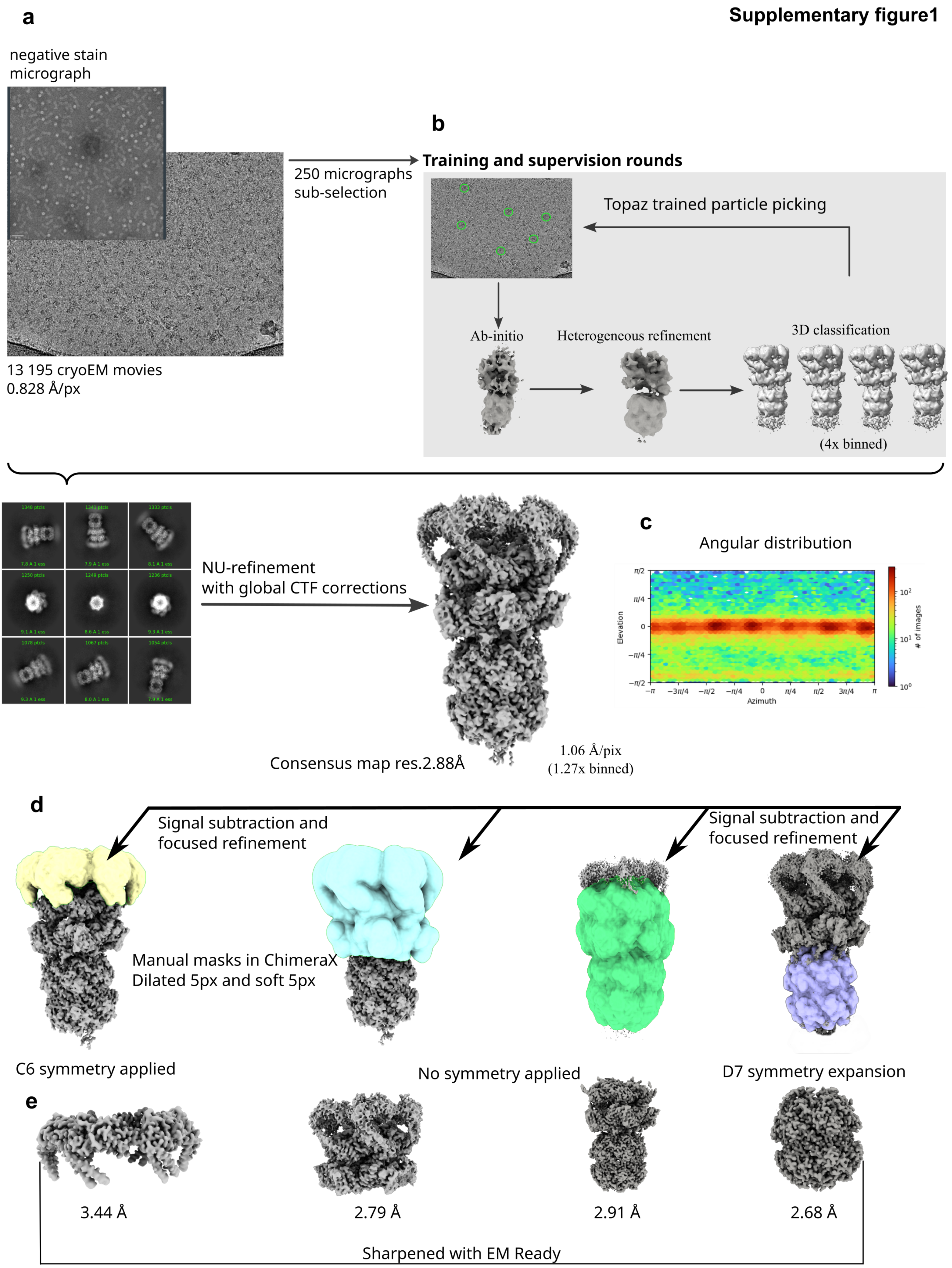

Supplementary figure1
a
negative stain
micrograph
250 micrographs
sub-selection
13 195 cryoEM movies
0.828 Å/px
b
Training and supervision rounds
Topaz trained particle picking
Ab-initio
Heterogeneous refinement
3D classification
(4x binned)
NU-refinement
with global CTF corrections
Consensus map res.2.88Å
1.06 Å/pix
(1.27x binned)
c
Angular distribution
Elevation
Azimuth
### of images
d
Signal subtraction and
focused refinement
Signal subtraction and
focused refinement
Manual masks in ChimeraX
Dilated 5px and soft 5px
C6 symmetry applied
No symmetry applied
D7 symmetry expansion
e
3.44 Å
2.79 Å
2.91 Å
2.68 Å
Sharpened with EM Ready

#### Supplementary Figure2

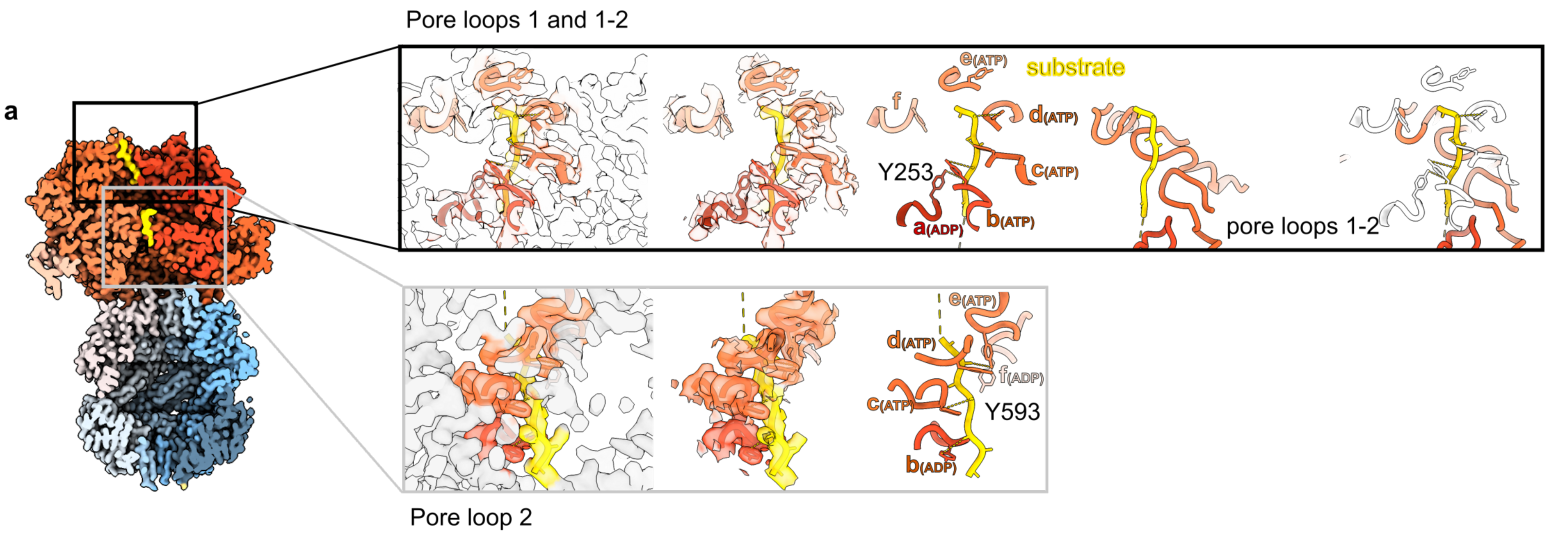

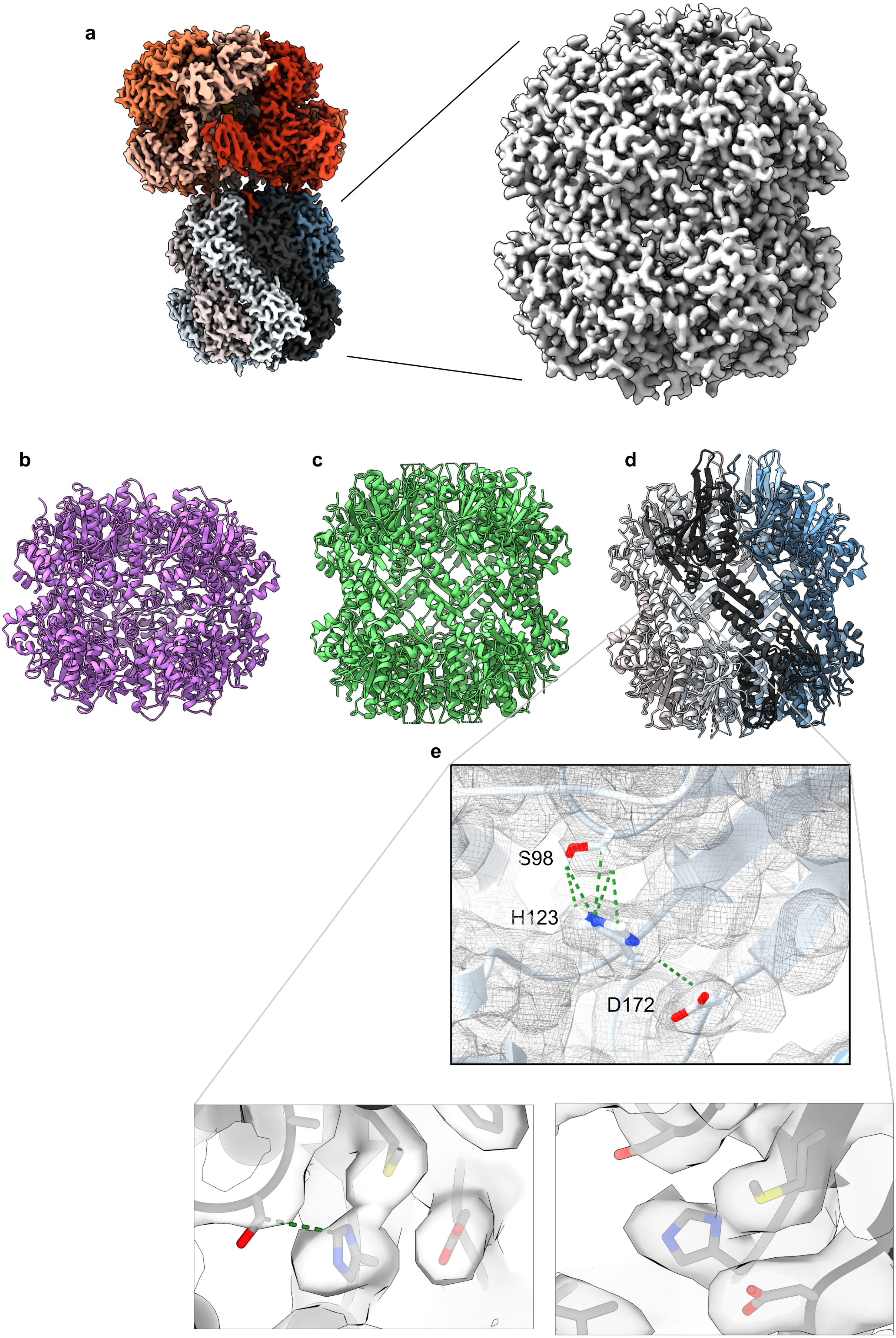


**Supplementary Figure3**

**Supplementary Figure4**

**a**

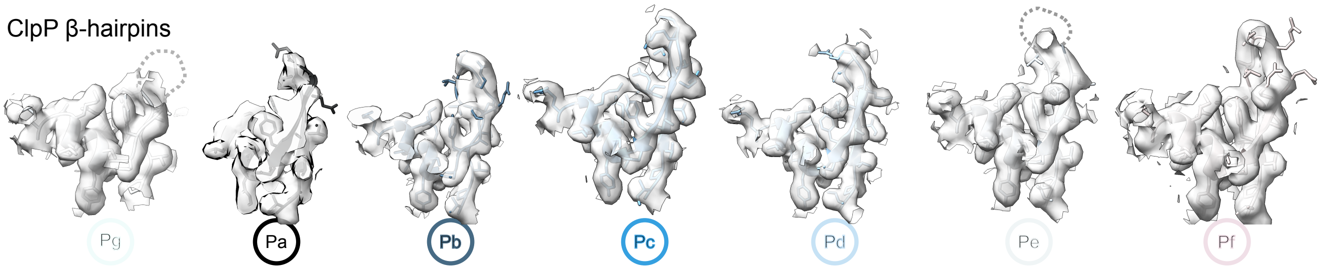


**b**

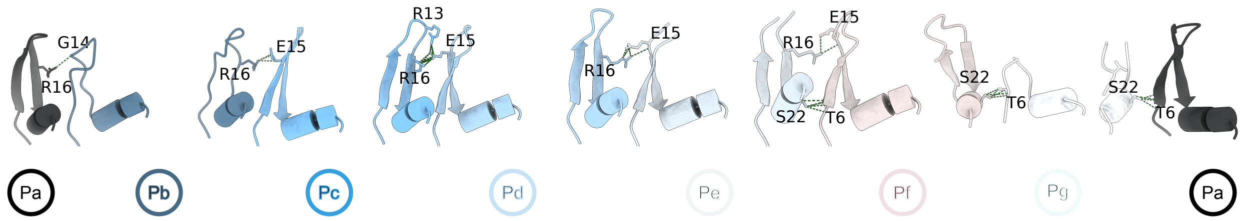

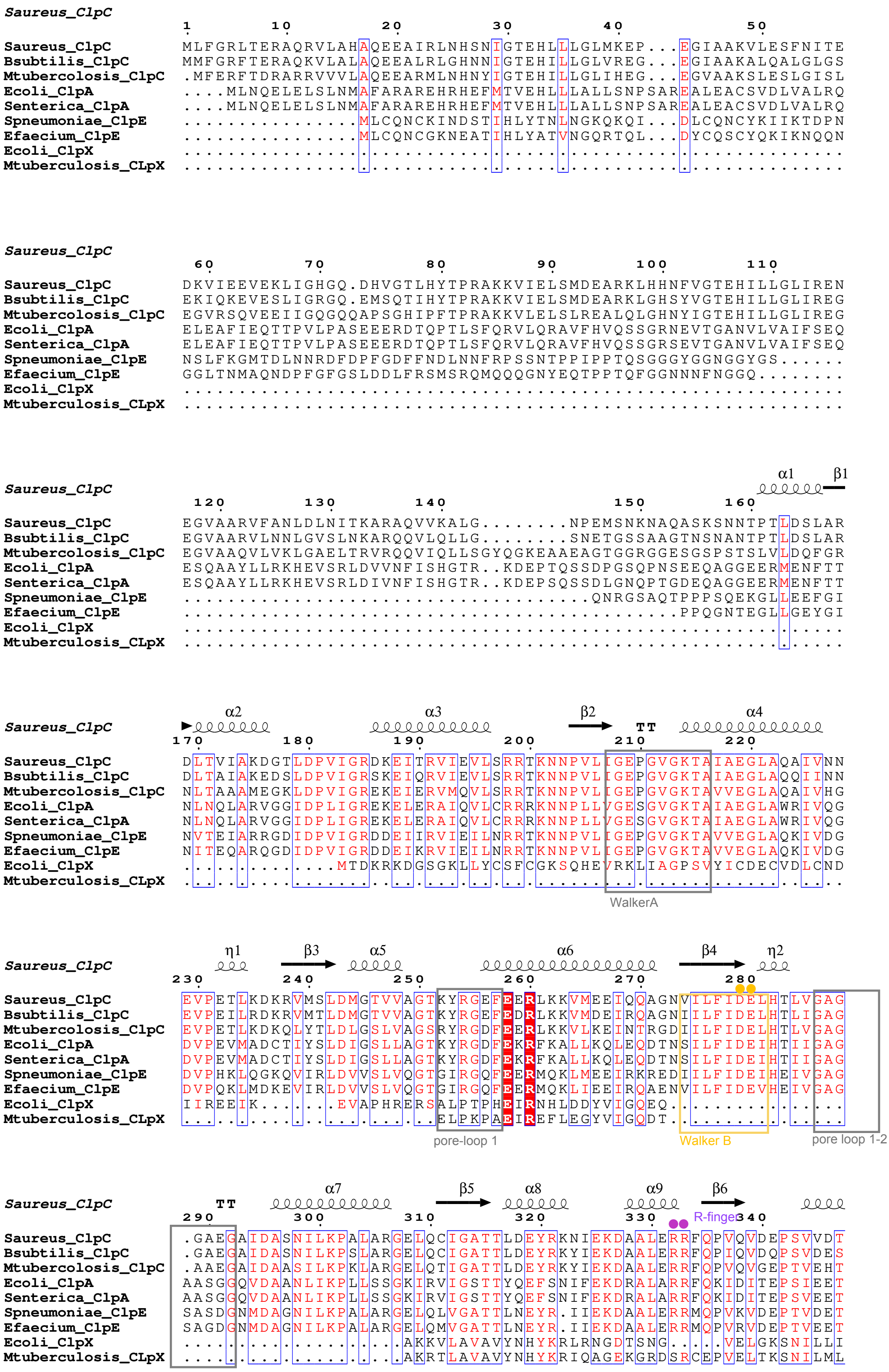
Saureus_ClpC
1 10 20 30 40 50
Saureus_ClpC MLFGRLTERAQRVLAHAQEEAIRLNHSNIGTEHLLLGLMKEP...EGIAAKVLESFNITE
Bsubtilis_ClpC MMFGRFTERAQKVLALAQEEALRLGHNNIGTEHILLGLVREG...EGIAAKALQALGLGS
Mtubercolosis_ClpC .MFERFTDRARRVVVLAQEEARMLNHNYIGTEHILLGLIHEG...EGVAAKSLESLGISL
Ecoli_ClpA ....MLNQELELSLNMAFARAREHRHEFMTVEHLLLALLSNPSAREALEACSVDLVALRQ
Senterica_ClpA ....MLNQELELSLNMAFARAREHRHEFMTVEHLLLALLSNPSAREALEACSVDLVALRQ
Spneumoniae_ClpE ...............MLCQNCKINDSTIHLYTNLNGKQKQI...DLCQNCYKIIKTDPN
Efaecium_ClpE ...............MLCQNCGKNEATIHLYATVNGQRTQL...DYCQSCYQKIKNQQN
Ecoli_ClpX ............................................................
Mtuberculosis_CLpX ............................................................
Saureus_ClpC
60 70 80 90 100 110
Saureus_ClpC DKVIEEVEKLIGHGQ.DHVGTLHYTPRAKKVIELSMDEARKLHHNFVGTEHILLGLIREN
Bsubtilis_ClpC EKIQKEVESLIGRGQ.EMSQTIHYTPRAKKVIELSMDEARKLGHSYVGTEHILLGLIREG
Mtubercolosis_ClpC EGVRSQVEEIIGQGQQAPSGHIPFTPRAKKVLELSLREALQLGHNYIGTEHILLGLIREG
Ecoli_ClpA ELEAFIEQTTPVLPASEEERDTQPTLSFQRVLQRAVFHVQSSGRNEVTGANVLVAIFSEQ
Senterica_ClpA ELEAFIEQTTPVLPASEEERDTQPTLSFQRVLQRAVFHVQSSGRSEVTGANVLVAIFSEQ
Spneumoniae_ClpE NSLFKGMTDLNNRDFDPFGDFFNDLNNFRPSSNTPPIPPTQSGGGYGGNGGYGS......
Efaecium_ClpE GGLTNMAQNDPFGFGSLDDLFRSMSRQMQQQGNYEQTPPTQFGGNNNFNGGQ........
Ecoli_ClpX ............................................................
Mtuberculosis_CLpX ............................................................
α1 β1
Saureus_ClpC
120 130 140 150 160
Saureus_ClpC EGVAARVFANLDLNITKARAQVVKALG........NPEMSNKNAQASKSNNTPTLDSLAR
Bsubtilis_ClpC EGVAARVLNNLGVSLNKARQQVLQLLG........SNETGSSAAGTNSNANTPTLDSLAR
Mtubercolosis_ClpC EGVAAQVLVKLGAELTRVRQQVIQLLSGYQGKEAAEAGTGGRGGESGSPSTSLVLDQFGR
Ecoli_ClpA ESQAAYLLRKHEVSRLDVVNFISHGTR..KDEPTQSSDPGSQPNSEEQAGGEERMENFTT
Senterica_ClpA ESQAAYLLRKHEVSRLDIVNFISHGTR..KDEPSQSSDLGNQPTGDEQAGGEERMENFTT
Spneumoniae_ClpE ....................................QNRGSAQTPPPSQEKGLLEEFGI
Efaecium_ClpE ...............................................PPQGNTEGLLGEYGI
Ecoli_ClpX ............................................................
Mtuberculosis_CLpX ............................................................
α2 α3 β2 TT α4
Saureus_ClpC
170 180 190 200 210 220
Saureus_ClpC DLTVIAKDGTLDPVIGRDKEITRVIEVLSRRTKNNPVLIGEPGVGKTAIAEGLAQAIVNN
Bsubtilis_ClpC DLTAIAKEDSLDPVIGRSKEIQRVIEVLSRRTKNNPVLIGEPGVGKTAIAEGLAQQIINN
Mtubercolosis_ClpC NLTAAAMEGKLDPVIGREKEIERVMQVLSRRTKNNPVLIGEPGVGKTAVVEGLAQAIVHG
Ecoli_ClpA NLNQLARVGGIDPLIGREKELERAIQVLCRRRKNNPLLVGESGVGKTAIAEGLAWRIVQG
Senterica_ClpA NLNQLARVGGIDPLIGREKELERAIQVLCRRRKNNPLLVGESGVGKTAIAEGLAWRIVQG
Spneumoniae_ClpE NVTEIARRGDIDPVIGRDDEIIRVIEILNRRTKNNPVLIGEPGVGKTAVVEGLAQKIVDG
Efaecium_ClpE NITEQARQGDIDPVIGRDDEIKRVIEILNRRTKNNPVLIGEPGVGKTAVVEGLAQKIVDG
Ecoli_ClpX ...............MTDKRKDGSGKLLYCSFCGKSQHEVRKLIAGPSVYICDECVDLCND
Mtuberculosis_CLpX ............................................................
WalkerA
η1 β3 α5 α6 β4 η2
Saureus_ClpC
230 240 250 260 270 280
Saureus_ClpC EVPETLKDKRVMSLDMGTVVAGTKYRGEFEERLKKVMEEIQQAGNVILFIDELHTLVGAG
Bsubtilis_ClpC EVPEILRDKRVMTLDMGTVVAGTKYRGEFEDRLKKVMDEIRQAGNIILFIDELHTLIGAG
Mtubercolosis_ClpC EVPETLKDKQLYTLDLGSLVAGSRYRGDFEERLKKVLKEINTRGDIILFIDELHTLVGAG
Ecoli_ClpA DVPEVMADCTIYSLDIGSLLAGTKYRGDFEKRFKALLKQLEQDTNSILFIDEIHTIIGAG
Senterica_ClpA DVPEVMADCTIYSLDIGSLLAGTKYRGDFEKRFKALLKQLEQDTNSILFIDEIHTIIGAG
Spneumoniae_ClpE DVPHKLQGKQVIRLDVVSLVQGTGIRGQFEERMQKLMEEIRKREDIILFIDEIHEIVGAG
Efaecium_ClpE DVPQKLMDKEVIRLDVVSLVQGTGIRGQFEERMQKLIEEIRQAENVILFIDEVHEIVGAG
Ecoli_ClpX IIREEIK......EVAPHRERSALPTPHEIRNHLDDYVIGQEQ.................
Mtuberculosis_CLpX ......................ELPKPAEIREFLEGYVIGQDT.................
pore-loop 1
Walker B
pore loop 1-2
TT α7 β5 α8 α9 β6
Saureus_ClpC
290 300 310 320 330 340
R-finger
Saureus_ClpC .GAEGAIDASNILKPALARGELQCIGATTLDEYRKNIEKDAALERRFQPVQVDEPSVVDT
Bsubtilis_ClpC .GAEGAIDASNILKPSLARGELQCIGATTLDEYRKYIEKDAALERRFQPIQVDQPSVDES
Mtubercolosis_ClpC .AAEGAIDAASILKPKLARGELQTIGATTLDEYRKYIEKDAALERRFQPVQVGEPTVEHT
Ecoli_ClpA AASGGQVDAANLIKPLLSSGKIRVIGSTTYQEFSNIFEKDRALARRFQKIDITEPSIEET
Senterica_ClpA AASGGQVDAANLIKPLLSSGKIRVIGSTTYQEFSNIFEKDRALARRFQKIDITEPSVEET
Spneumoniae_ClpE SASDGNMDAGNILKPALARGELQLVGATTLNEYR.IIEKDAALERRMQPVKVDEPTVDET
Efaecium_ClpE SAGDGNMDAGNILKPALARGELQMVGATTLNEYR.IIEKDAALERRMQPVRVDEPTVEET
Ecoli_ClpX ....................AKKVLAVAVYNHYKRLRNGDTSNG.....VELGKSNILLI
Mtuberculosis_CLpX ....................AKRTLAVAVYNHYKRIQAGEKGRDSRCEPVELTKSNILML

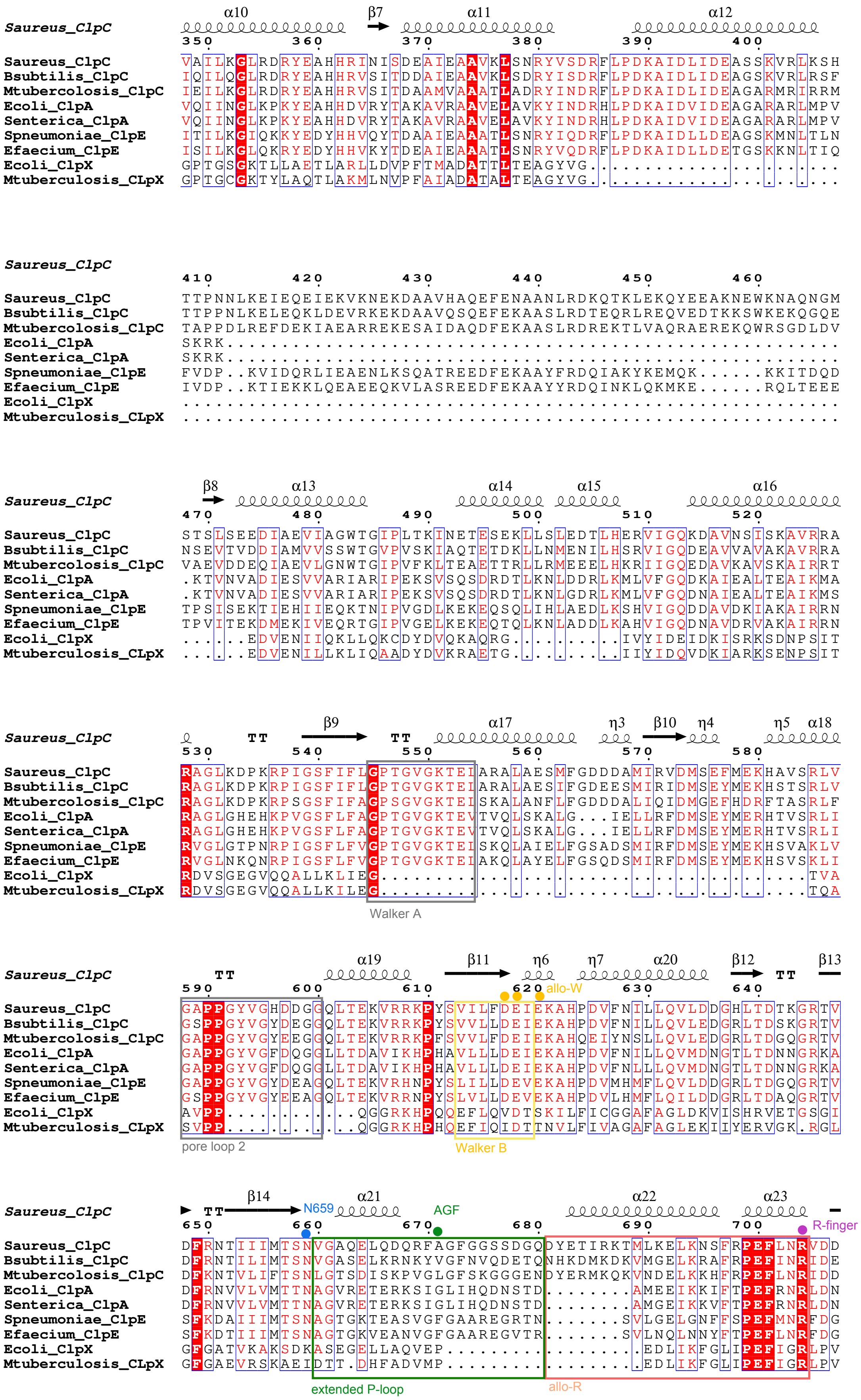

Saureus_ClpC
α10 β7 α11 α12
350 360 370 380 390 400
Saureus_ClpC VAILKGLRDRYEAHHRINISDEAIEAAVKLSNRYVSDRFLPDKAIDLIDEASSKVRLKSH
Bsubtilis_ClpC IQILQGLRDRYEAHHRVSITDDAIEAAVKLSDRYISDRFLPDKAIDLIDEAGSKVRLRSF
Mtubercolosis_ClpC IEILKGLRDRYEAHHRVSITDAAMVAAATLADRYINDRFLPDKAIDLIDEAGARMRIRRM
Ecoli_ClpA VQIINGLKPKYEAHHDVRYTAKAVRAAVELAVKYINDRHLPDKAIDVIDEAGARARLMPV
Senterica_ClpA VQIINGLKPKYEAHHDVRYTAKAVRAAVELAVKYINDRHLPDKAIDVIDEAGARARLMPV
Spneumoniae_ClpE ITILKGIQKKYEDYHHVQYTDAAIEAAATLSNRYIQDRFLPDKAIDLLDEAGSKMNLTLN
Efaecium_ClpE ISILKGLQKRYEDYHHVKYTDEAIEAAATLSNRYVQDRFLPDKAIDLLDETGSKKNLTIQ
Ecoli_ClpX GPTGSGKTLLAETLARLLDVPFTMADATTLTEAGYVG.......................
Mtuberculosis_CLpX GPTGCGKTYLAQTLAKMLNVPFAIADATALTEAGYVG.......................
Saureus_ClpC
410 420 430 440 450 460
Saureus_ClpC TTPNNLKEIEQEIEKVKNEKDAAVHAQEFENAANLRDKQTKLEKQYEEAKNEWKNAQNGM
Bsubtilis_ClpC TTPPNLKELEQKLDEVRKEKDAAVQSQEFEKAASLRDTEQRLREQVEDTKKSWKEKQGQE
Mtubercolosis_ClpC TAPPDLREFDEKIAEARREKESAIDAQDFEKAASLRDREKTLVAQRAEREKQWRSGDLDV
Ecoli_ClpA SKRK........................................................
Senterica_ClpA SKRK........................................................
Spneumoniae_ClpE FVDP..KVIDQRLIEAENLKSQATREEDFEKAAYFRDQIAKYKEMQK......KKITDQD
Efaecium_ClpE IVDP..KTIEKKLQEAEEQKVLASREEDFEKAAYYRDQINKLQKMKE......RQLTEEE
Ecoli_ClpX ............................................................
Mtuberculosis_CLpX ............................................................
Saureus_ClpC
β8 α13 α14 α15 α16
470 480 490 500 510 520
Saureus_ClpC STSLSEEDIAEVIAGWTGIPLTKINETESEKLLSLEDTLHERVIGQKDAVNSISKAVRRA
Bsubtilis_ClpC NSEVTVDDIAMVVSSWTGVPVSKIAQTETDKLLNMENILHSRVIGQDEAVVAVAKAVRRA
Mtubercolosis_ClpC VAEVDDEQIAEVLGNWTGIPVFKLTEAETTRLLRMEEELHKRIIGQEDAVKAVSKAIRRT
Ecoli_ClpA .KTVNVADIESVVARIARIPEKSVSQSDRDTLKNLDDRLKMLVFGQDKAIEALTEAIKMA
Senterica_ClpA .KTVNVADIESVVARIARIPEKSVSQSDRDTLKNLGDRLKMLVFGQDNAIEALTEAIKMS
Spneumoniae_ClpE TPSISEKTIEHIIEQKTNIPVGDLKEKEQSQLIHLAEDLKSHVIGQDDAVDKIAKAIRRN
Efaecium_ClpE TPVITEKDMEKIVEQRTGIPVGELKEKEQTQLKNLADDLKAHVIGQDNAVDRVAKAIRRN
Ecoli_ClpX ......EDVENIIQKLLQKCDYDVQKAQRG..........IVYIDEIDKISRKSDNPSIT
Mtuberculosis_CLpX ......EDVENILLKLIQAADYDVKRAETG..........IIYIDQVDKIARKSENPSIT
Saureus_ClpC
TT β9 TT α17 η3 β10 η4 η5 α18
530 540 550 560 570 580
Saureus_ClpC RAGLKDPKRPIGSFIFLGPTGVGKTEIARALAESMFGDDDAMIRVDMSEFMEKHAVSRLV
Bsubtilis_ClpC RAGLKDPKRPIGSFIFLGPTGVGKTEIARALAESIFGDEESMIRIDMSEYMEKHSTSRLV
Mtubercolosis_ClpC RAGLKDPKRPSGSFIFAGPSGVGKTEISKALANFLFGDDDALIQIDMGEFHDRFTASRLF
Ecoli_ClpA RAGLGHEHKPVGSFLFAGPTGVGKTEVTVQLSKALG...IELLRFDMSEYMERHTVSRLI
Senterica_ClpA RAGLGHEHKPVGSFLFAGPTGVGKTEVTVQLSKALG...IELLRFDMSEYMERHTVSRLI
Spneumoniae_ClpE RVGLGTPNRPIGSFLFVGPTGVGKTEISKQLAIELFGSADSMIRFDMSEYMEKHSVAKLV
Efaecium_ClpE RVGLNKQNRPIGSFLFVGPTGVGKTEIAKQLAYELFGSQDSMIRFDMSEYMEKHSVSKLI
Ecoli_ClpX RDVSGEGVQQALLKLIEG.......................................TVA
Mtuberculosis_CLpX RDVSGEGVQQALLKILEG.......................................TQA
Walker A
Saureus_ClpC
TT α19 β11 η6 η7 α20 β12 TT β13
590 600 610 620 allo-W 630 640
Saureus_ClpC GAPPGYVGHDDGGQLTEKVRRKPYSVILFDEIEKAHPDVFNILLQVLDDGHLTDTKGRTV
Bsubtilis_ClpC GSPPGYVGYDEGGQLTEKVRRKPYSVVLLDEIEKAHPDVFNILLQVLEDGRLTDSKGRTV
Mtubercolosis_ClpC GAPPGYVGYEEGGQLTEKVRRKPFSVVLFDEIEKAHQEIYNSLLQVLEDGRLTDGQGRTV
Ecoli_ClpA GAPPGYVGFDQGGLLTDAVIKHPHAVLLLDEIEKAHPDVFNILLQVMDNGTLTDNNGRKA
Senterica_ClpA GAPPGYVGFDQGGLLTDAVIKHPHAVLLLDEIEKAHPDVFNLLLQVMDNGTLTDNNGRKA
Spneumoniae_ClpE GAPPGYVGYDEAGQLTEKVRHNPYSLILLDEVEKAHPDVMHMFLQVLDDGRLTDGQGRTV
Efaecium_ClpE GSPPGYVGYEEAGQLTEKVRRNPYSLVLLDEVEKAHPDVLHMFLQILDDGRLTDAQGRTV
Ecoli_ClpX AVPP............QGGRKHPQQEFLQVDTSKILFICGGAFAGLDKVISHRVETGSGI
Mtuberculosis_CLpX SVPP............QGGRKHPHQEFIQIDTTNVLFIVAGAFAGLEKIIYERVGK.RGL
pore loop 2
Walker B
Saureus_ClpC
TT β14 N659 α21 AGF α22 α23
650 660 670 680 690 700 R-finger
Saureus_ClpC DFRNTIIIMTSNVGAQELQDQRFAGFGGSSDGQDYETIRKTMLKELKNSFRPEFLNRVDD
Bsubtilis_ClpC DFRNTILIMTSNVGASELKRNKYVGFNVQDETQNHKDMKDKVMGELKRAFRPEFINRIDE
Mtubercolosis_ClpC DFKNTVLIFTSNLGTSDISKPVGLGFSKGGGENDYERMKQKVNDELKKHFRPEFLNRIDD
Ecoli_ClpA DFRNVVLVMTTNAGVRETERKSIGLIHQDNSTD........AMEEIKKIFTPEFRNRLDN
Senterica_ClpA DFRNVVLVMTTNAGVRETERKSIGLIHQDNSTD........AMGEIKKVFTPEFRNRLDN
Spneumoniae_ClpE SFKDAIIIMTSNAGTGKTEASVGFGAAREGRTN.......SVLGELGNFFSPEFMNRFDG
Efaecium_ClpE SFKDTIIIMTSNAGTGKVEANVGFGAAREGVTR.......SVLNQLNNYFTPEFLNRFDG
Ecoli_ClpX GFGATVKAKSDKASEGELLAQVEP..................EDLIKFGLIPEFIGRLPV
Mtuberculosis_CLpX GFGAEVRSKAEIDTT.DHFADVMP..................EDLIKFGLIPEFIGRLPV
extended P-loop
allo-R

```
                      β15           α24                 η8   β16     α25
Saureus_ClpC
                      710       720       730       740       750       760
Saureus_ClpC          IIVFHKLTKEELKEIVTMMVNKLTNRLSEQNINIIVTDKAKDKIAEEGYDPEYGARPLIR
Bsubtilis_ClpC        IIVFHSLEKKHLTEIVSLMSDQLTKRLKEQDLSIELTDAAKAKVAEEGVDLEYGARPLRR
Mtubercolosis_ClpC    IIVFHQLTREEIIRMVDLMISRVAGQLKSKDMALVLTDAAKALLAKRGFDPVLGARPLRR
Ecoli_ClpA            IIWFDHLSTDVIHQVVDKFIVELQVQLDQKGVSLEVSQEARNWLAEKGYDRAMGARPMAR
Senterica_ClpA        IIWFDHLSGEVIHQVVDKFIVELQAQLDQKGVSLEVSQEARDWLAEKGYDRAMGARPMAR
Spneumoniae_ClpE      IIEFKALSKDNLLQIVELMLADVNKRLSSNNIRLDVTDKVKEKLVDLGYDPKMGARPLRR
Efaecium_ClpE         IIEFSALSKENLMTIVTLMLDDVNQMLAAQQLHIEVPTNVKEKLVDLGYDPSMGARPLRR
Ecoli_ClpX            VATLNELSEEALIQILKEPKNALTKQYQALFNLEGVDLEFRDEALDAIAKKAMARKTGAR
Mtuberculosis_CLpX    VASVTNLDKESLVKILSEPKNALVKQYIRLFEMDGVELEFTDDALEAIADQAIHRGTGAR

                        α26                     β17     TT   β18
Saureus_ClpC
                      770       780       790       800       810
Saureus_ClpC          AIQKTIEDNLSELILDGNQIEGKKVTVDHDGKEFKYDIAEQTSETKTPSQV.........
Bsubtilis_ClpC        AIQKHVEDRLSEELLRGNIHKGQHIVLDVEDGEFVVKTTAKTN.................
Mtubercolosis_ClpC    TIQREIEDQLSEKILFEEVGPGQVVTVDVDNWDGEGPGEDAVFTFTGTRKPPAEPDLAKA
Ecoli_ClpA            VIQDNLKKPLANELLFGSLVDGGQVTVALDKEKNELTYGFQSAQKHKAEAAH........
Senterica_ClpA        VIQDNLKKPLANELLFGSLVDGGQVTVALDKEKNALTYGFQSAQKHKPEAAH........
Spneumoniae_ClpE      TIQDYIEDTITDYYLENPSEKDLKAVMTSKGNIQIKSAKKAEVKSSEKEK..........
Efaecium_ClpE         TIQEQIEDGIAEFYLDHPSIHELKAKLDKDGKIIVTSKPERLAKESAEETAE........
Ecoli_ClpX            GLRSIVEAALLDTMYDLPSMEDVEKVVIDESVIDGQSKPLLIYGKPEAQQASGE......
Mtuberculosis_CLpX    GLRAIMEEVLLPVMYDIPSRDDVAKVVVTKETVQDNVLPTIVPR................

Saureus_ClpC

Saureus_ClpC          .............
Bsubtilis_ClpC        .............
Mtubercolosis_ClpC    GAHSAGGPEPAAR
Ecoli_ClpA            .............
Senterica_ClpA        .............
Spneumoniae_ClpE      .............
Efaecium_ClpE         .............
Ecoli_ClpX            .............
Mtuberculosis_CLpX    .............
```

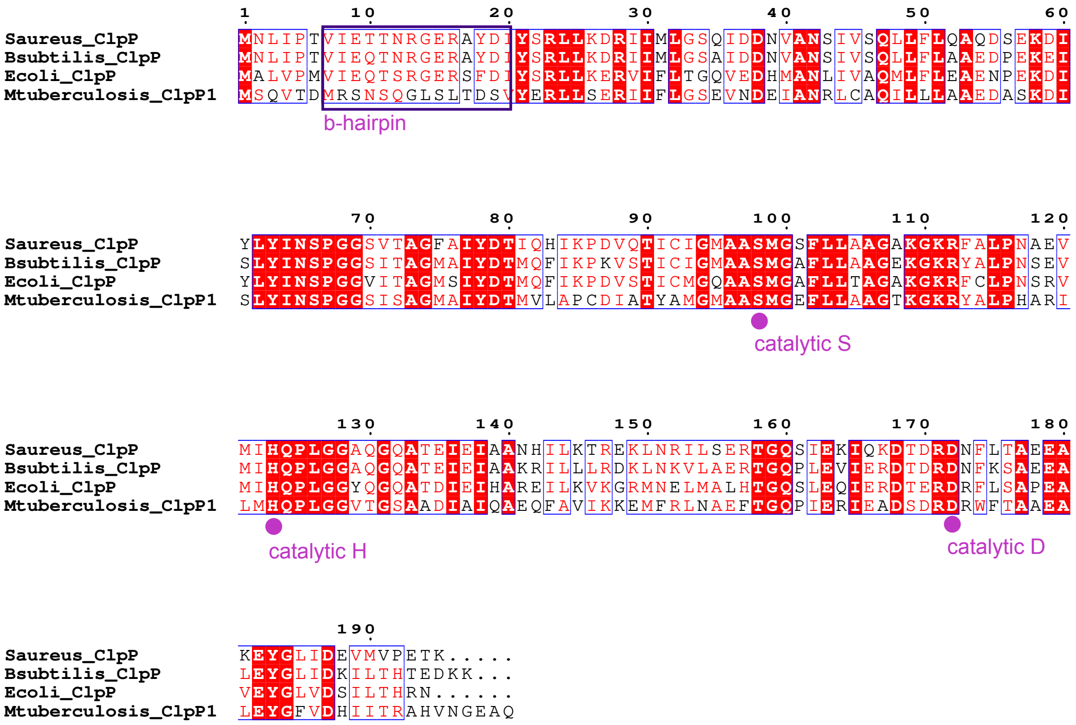
1 10 20 30 40 50 60
Saureus_ClpP MNLIPTVIETTNRGERAYDIYSRLLKDRIIMLGSQIDDNVANSIVSQLLFLQAQDSEKDI
Bsubtilis_ClpP MNLIPTVIEQTNRGERAYDIYSRLLKDRIIMLGSAIDDNVANSIVSQLLFLAAEDPEKEI
Ecoli_ClpP MALVPMVIEQTSRGERSFDIYSRLLKERVIFLTGQVEDHMANLIVAQMLFLEAENPEKDI
Mtuberculosis_ClpP1 MSQVTDMRSNSQGLSLTDSVYERLLSERIIFLGSEVNDEIANRLCAQILLLAAEDASKDI
b-hairpin
70 80 90 100 110 120
Saureus_ClpP YLYINSPGGSVTAGFAIYDTIQHIKPDVQTICIGMAASMGSFLLAAGAKGKRFALPNAEV
Bsubtilis_ClpP SLYINSPGGSITAGMAIYDTMQFIKPKVSTICIGMAASMGAFLLAAGEKGKRYALPNSEV
Ecoli_ClpP YLYINSPGGVITAGMSIYDTMQFIKPDVSTICMGQAASMGAFLLTAGAKGKRFCLPNSRV
Mtuberculosis_ClpP1 SLYINSPGGSISAGMAIYDTMVLAPCDIATYAMGMAASMGEFLLAAGTKGKRYALPHARI
catalytic S
130 140 150 160 170 180
Saureus_ClpP MIHQPLGGAQGQATEIEIAANHILKTREKLNRILSERTGQSIEKIQKDTDRDNFLTAEEA
Bsubtilis_ClpP MIHQPLGGAQGQATEIEIAAKRILLLRDKLNKVLAERTGQPLEVIERDTDRDNFKSAEEA
Ecoli_ClpP MIHQPLGGYQGQATDIEIHAREILKVKGRMNELMALHTGQSLEQIERDTERDRFLSAPEA
Mtuberculosis_ClpP1 LMHQPLGGVTGSAADIAIQAEQFAVIKKEMFRLNAEFTGQPIERIEADSDRDRWFTAAEA
catalytic H
catalytic D
190
Saureus_ClpP KEYGLIDEVMVPETK.....
Bsubtilis_ClpP LEYGLIDKILTHTEDKK...
Ecoli_ClpP VEYGLVDSILTHRN......
Mtuberculosis_ClpP1 LEYGFVDHIITRAHVNGEAQ

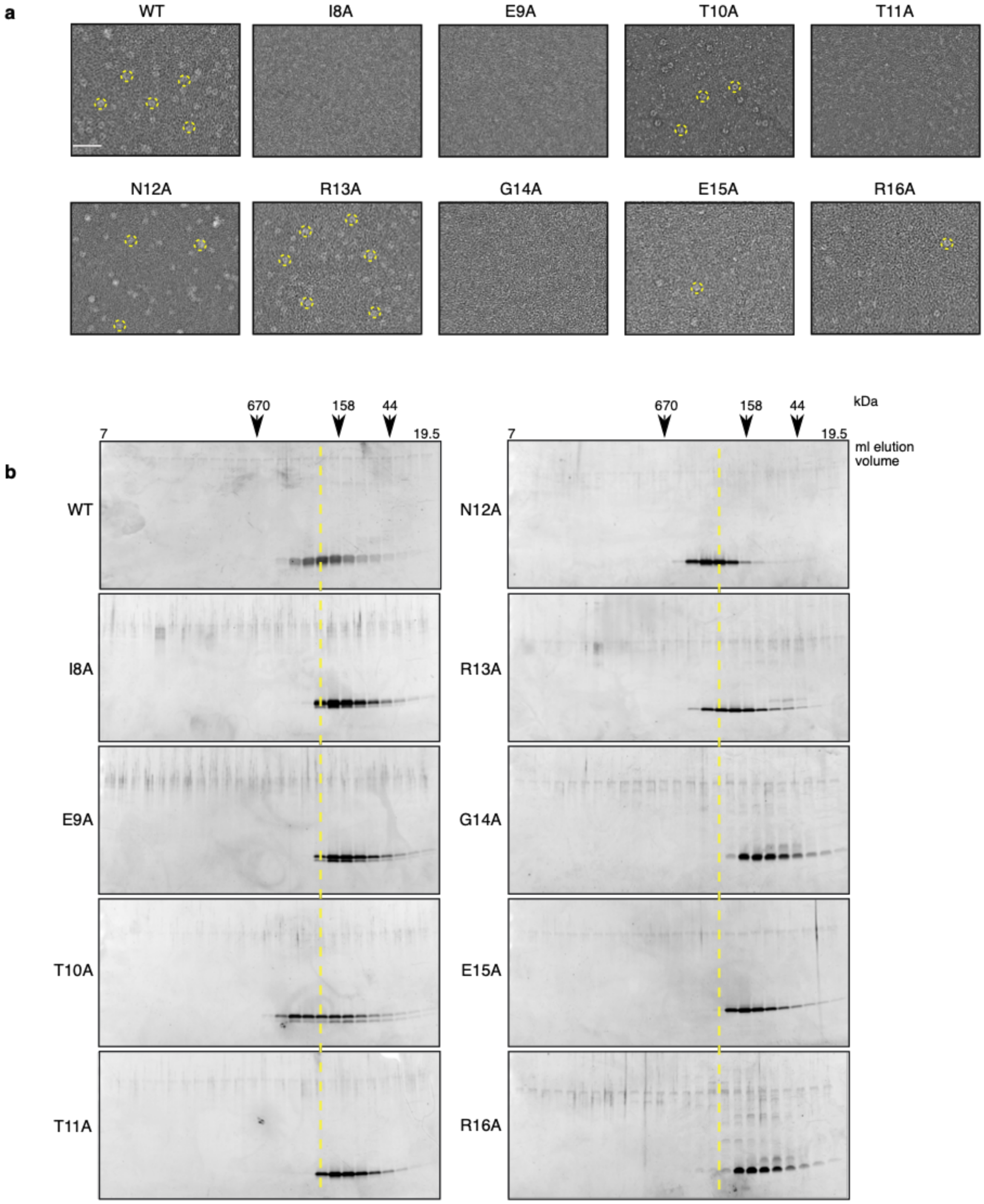
a
WT
I8A
E9A
T10A
T11A
N12A
R13A
G14A
E15A
R16A
b
670
158
44
kDa
7
19.5
ml elution volume
WT
I8A
E9A
T10A
T11A
N12A
R13A
G14A
E15A
R16A

**Supplementary Figure 7**

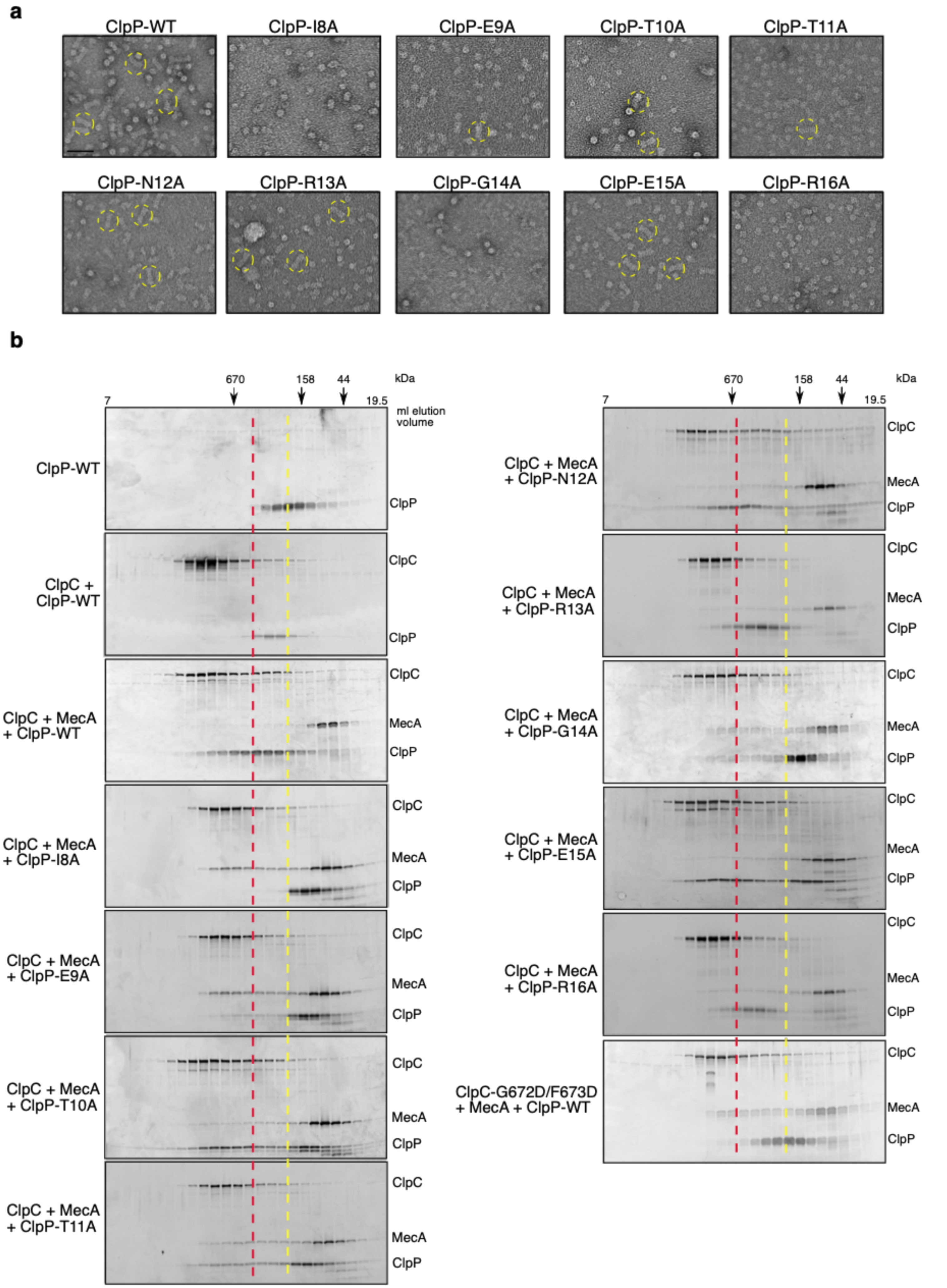

**a**

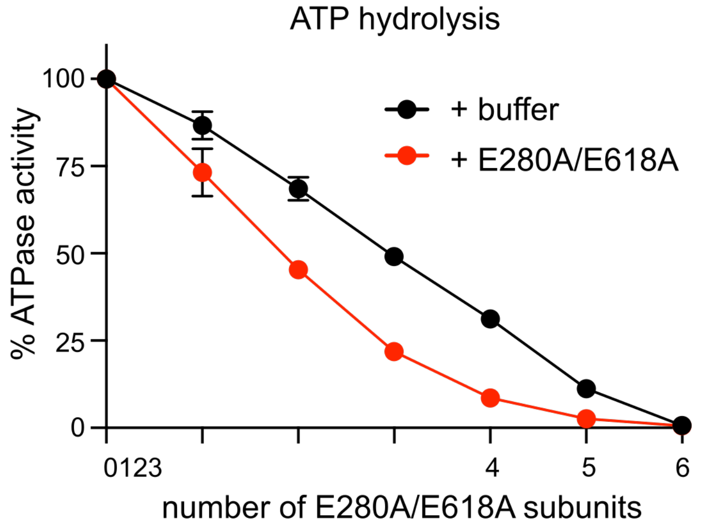
ATP hydrolysis
% ATPase activity
100
75
50
25
0
+ buffer
+ E280A/E618A
0123
4
5
6
number of E280A/E618A subunits

**b**

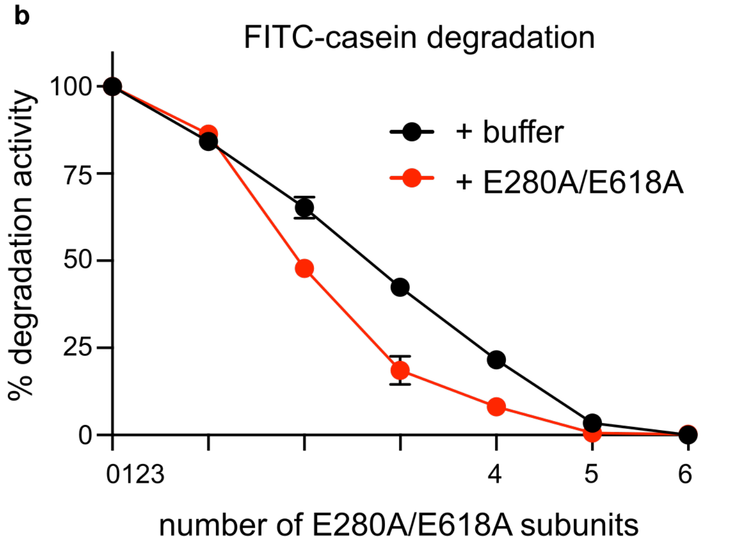
FITC-casein degradation
% degradation activity
100
75
50
25
0
+ buffer
+ E280A/E618A
0123
4
5
6
number of E280A/E618A subunits

**c**

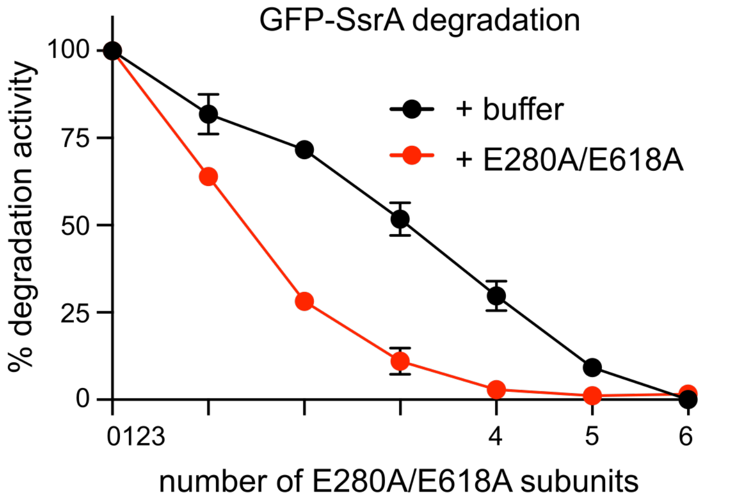
GFP-SsrA degradation
% degradation activity
100
75
50
25
0
+ buffer
+ E280A/E618A
0123
4
5
6
number of E280A/E618A subunits

**d**

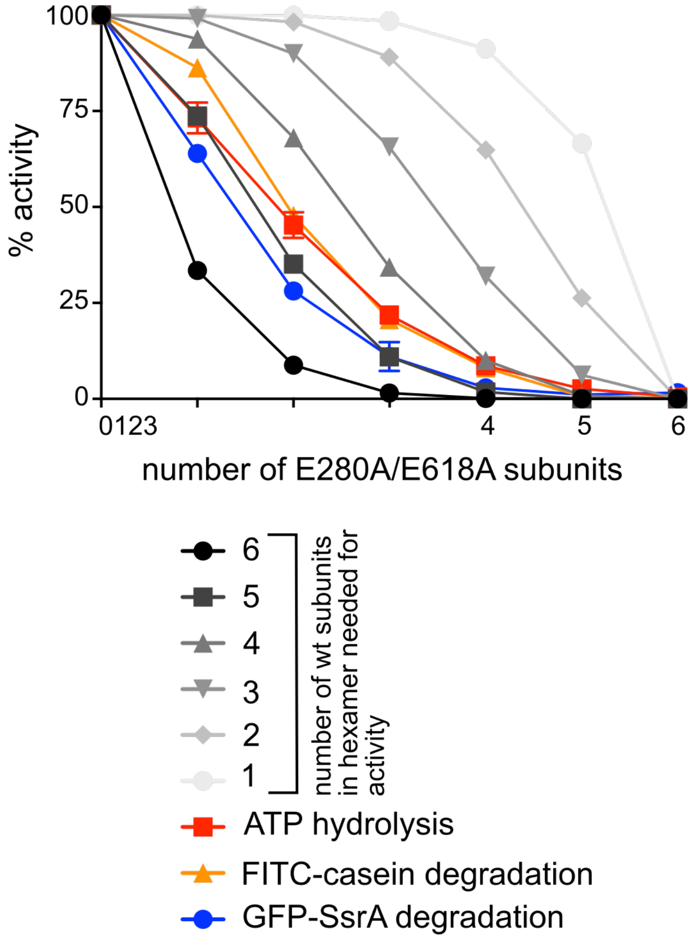
% activity
100
75
50
25
0
0123
4
5
6
number of E280A/E618A subunits
6
5
4
3
2
1
number of wt subunits in hexamer needed for activity
ATP hydrolysis
FITC-casein degradation
GFP-SsrA degradation

**Supplementary Figure9**

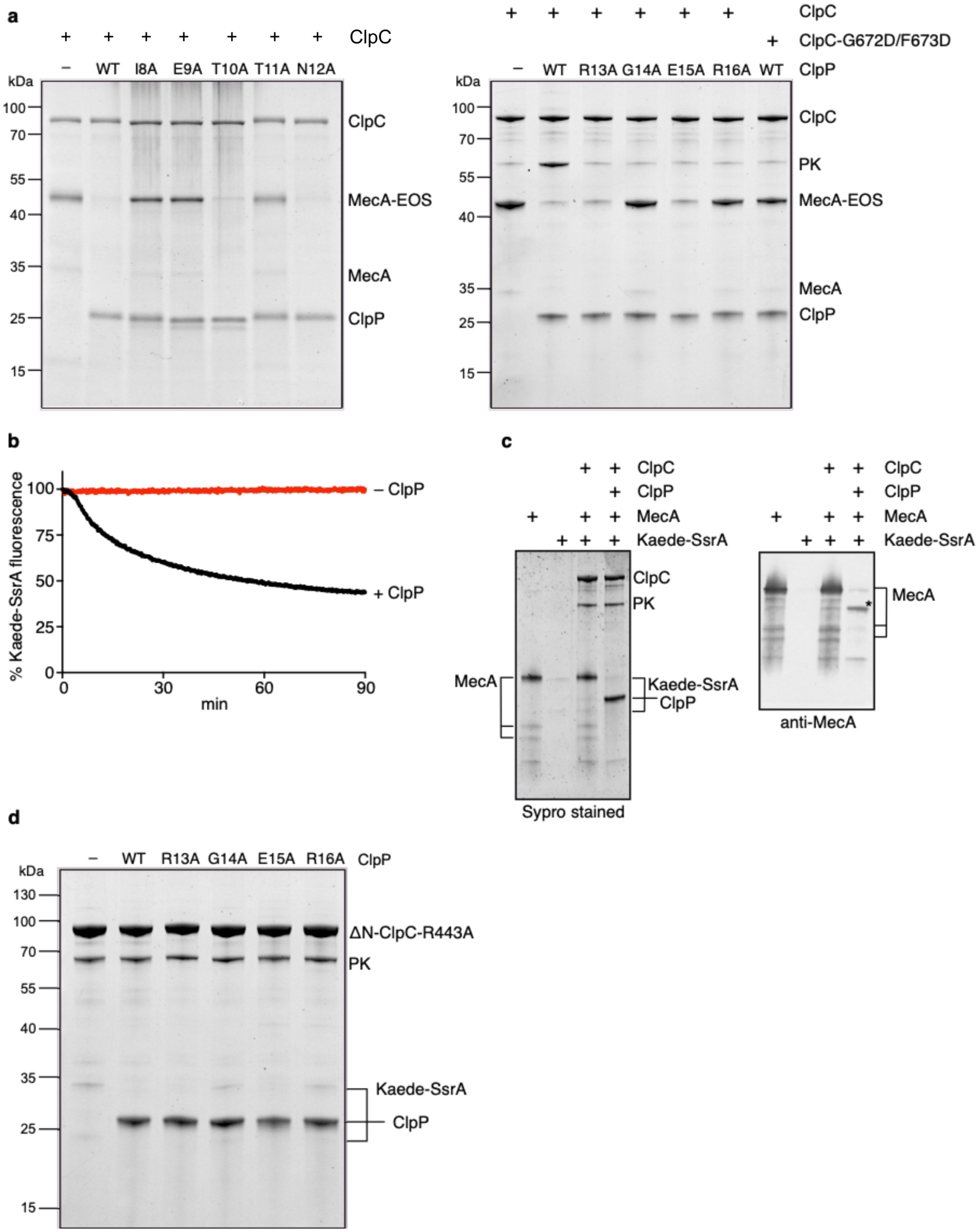

**Supplementary figure 10**

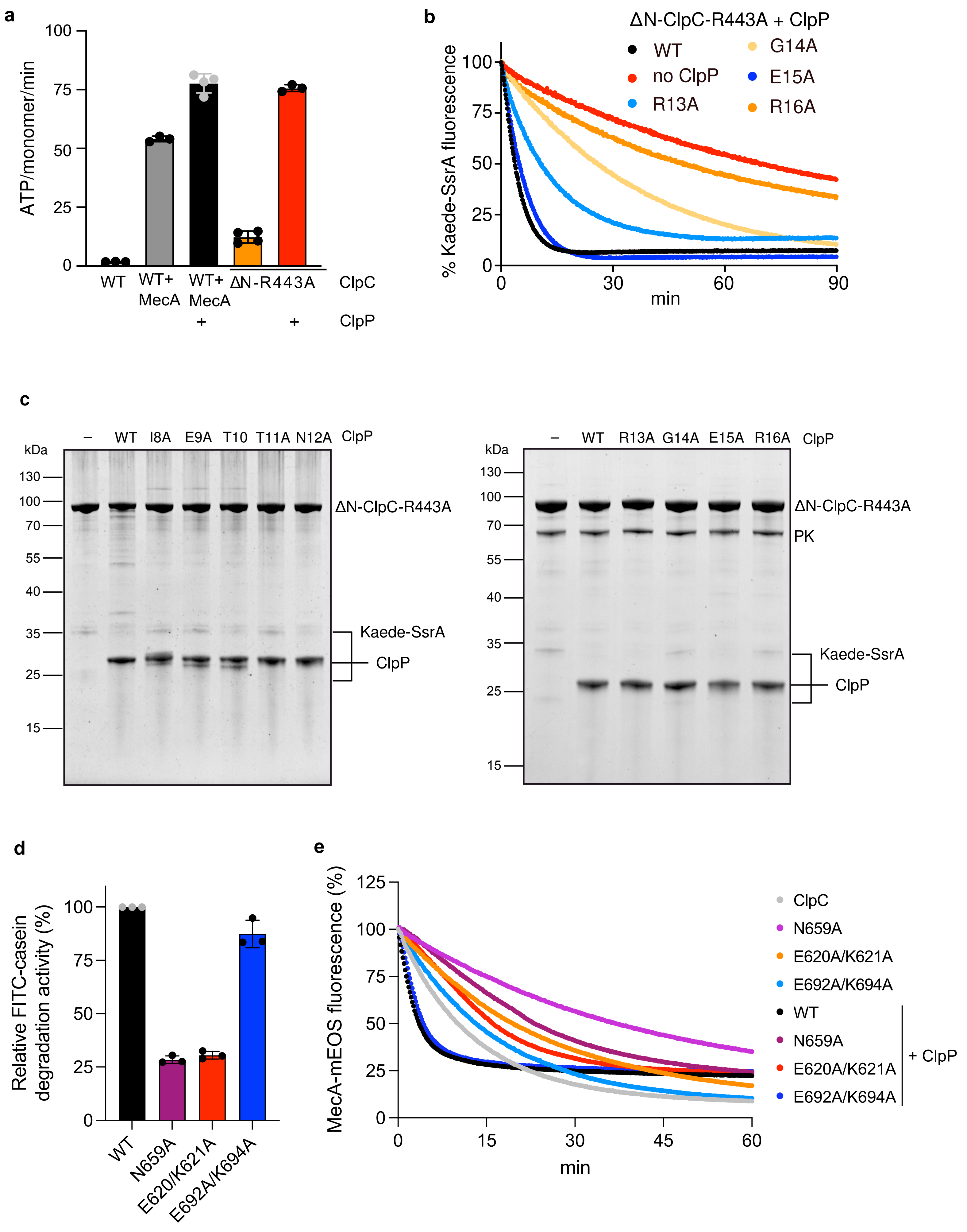

*Supplementary Table 1:* ***Characterization of S. aureus ClpP β-hairpins mutants.***

| ClpP variant | ClpP 14-mer formation | | MecA/ClpC/ClpP complex formation | | ClpP proteolytic activity | | | MecA/ClpC/ClpP proteolytic activity | | ClpC ATPase activation by ClpP | | Enhancement of ClpC unfolding activity by ClpP | |
|---|---|---|---|---|---|---|---|---|---|---|---|---|---|
| | EM | SEC | EM | SEC | LY-AMC | LY-AMC + ADEP1 | FITC-casein + ADEP1 | FITC-casein | GFP-SsrA | ClpC/MecA | ΔN-ClpC-R443A | ClpC/MecA-mEOS3.2 | ΔN-ClpC-R443A/Kaede-SsrA |
| WT | ++ | ++ | ++ | ++ | ++ | ++ | ++ | ++ | ++ | ++ | ++ | ++ | ++ |
| I8A | – | – | – | – | – | (+) | + | – | – | – | – | – | – |
| E9A | – | – | (+) | – | – | (+) | +++ | – | – | – | – | – | – |
| T10A | ++ | ++ | + | ++ | (+) | ++ | +++ | ++ | +(+) | ++ | +(+) | +(+) | +(+) |
| T11A | – | – | (+) | (+) | – | + | +++ | – | (+) | – | – | – | – |
| N12A | ++ | ++ | ++ | ++ | ++ | ++ | ++ | ++ | ++ | ++ | +(+) | ++ | ++ |
| R13A | ++ | + | ++ | + | ++ | ++ | + | + | + | + | + | + | + |
| G14A | – | – | – | – | – | ++ | +++ | – | – | – | – | – | – |
| E15A | (+) | – | ++ | ++ | – | ++ | +++ | (+) | + | (+) | + | (+) | +(+) |
| R16A | (+) | – | – | – | (+) | ++ | + | (+) | (+) | – | – | – | – |

Oligomerization and MecA/ClpC complex formation properties and proteolytic activities of ClpP β-hairpins mutants were qualitatively assessed. The properties and activities of ClpP-WT were defined as "++". ClpP mutants were categorized into three groups: no major defects (green), partial defects (orange) and severe defects (red).

**Supplementary table 2.** Cryo-EM data collection, refinement and validation statistics

| **Parameter** | **WT ClpC/ClpP body ClpC w/o NTD and MD (C1)**<br><br>**EMDB:** EMD-51367<br>**PDB:** 9GI1 | **WT MecA crown with ClpC NTD and MD (C6)**<br><br>**EMDB:** EMD-51498<br>**PDB:** 9GOQ | **WT clpP**<br><br>**(D7)**<br><br>**EMDB:** EMD-53538<br>**PDB:** 9R2S | **WT MecA/ClpC/ClpP Composite map**<br><br>**EMDB:** EMD-53879<br>**PDB:** 9RAI |
|---|---|---|---|---|
| Microscope | Titan Krios G3i (TFS) | Titan Krios G3i (TFS) | Titan Krios G3i (TFS) | Titan Krios G3i (TFS) |
| Detector and energy filter | Gatan K3 + Bioquantum(Ametek ) | Gatan K3 + Bioquantum(Ametek) | Gatan K3 + Bioquantum(Am etek) | Gatan K3 + Bioquantum(Ametek) |
| Nominal magnification (nominal/calibrated at detector) | 105k | 105k | 105k | 105k |
| Voltage (kV) | 300 | 300 | 300 | 300 |
| Defocus range (um) | -0.6 to -2.2 | -0.6 to -2.2 | -0.6 to -2.2 | -0.6 to -2.2 |
| Total electron exposure (or fluence, e-/Å2 | 40 | 40 | 40 | 40 |
| Exposure rate (or flux, e-/pixel/s) | 15 | 15 | 15 | 15 |
| Number of frames collected | 40 | 40 | 40 | 40 |
| Pixel size (Å) | 0.828 (binned final 1.059) | 0.828 (binned final 1.059) | 0.828 (binned final 1.059) | 0.994 |
| Energy filter slit width (eV) | 20 | 20 | 20 | 20 |
| Automation software | EPU v3.7 | EPU v3.7 | EPU v3.7 | EPU v3.7 |
| # Micrographs used | 12 990 | 12 990 | 12 990 | 12 990 |
| Total # of extracted particles | 2 851 121 | 2 851 121 | 2 851 121 | 2 851 121 |
| Total # of refined particles (particles after removing junk) | 82 854 | 82 854 | 82 854 | 82 854 |
| # of particles in final map | 48 011 | 82 854 | 82 854 | 82 854 |
| Resolution of unmasked and masked reconstructions at 0.143 FSC | 4.1/2.9 | 4.0/3.4 | 2.8/2.4 | N/A |
| Local resolution range (Å) | 2 to 6 | 2.3 to 5.8 | 2.3 to 3.4 | N/A |
| Map sharpening B factor (Å2) / (B factor Range) | -43 | -127 | -66 | N/A |
| Model composition<br>Non-hydrogen atoms<br>Protein residues | <br>47097<br>6024 | <br>13878<br>1704 | <br>19981<br>2591 | <br>60975<br>7728 |

| | | | | |
|---|---|---|---|---|
| Ligands | MG:7, AGS:6;ADP:4 | 0 | 0 | MG:7, AGS:6;ADP:4 |
| *B* factors (Å$^2$) (min/max/mean)<br>Protein<br>Ligand | 11.7/299.27/127.75<br>79.98/179.82/108.55 | 103.72/307.67/174.42 | 17.59/187.99/35.69 | 16.58/420.03/83.70<br>37.85/119.65/74.77 |
| Map sharpening EMReady (any) | Yes | Yes | Yes | Yes |
| Atomic modeling refinement package(s) | Phenix 1.21<br>Coot 0.95<br>Isolde | Phenix 1.21<br>Coot 0.95<br>Isolde | Phenix 1.21<br>Coot 0.95<br>Isolde | Phenix 1.21<br>Coot 0.95<br>Isolde |
| CCvolume/CCmask | 0.80 | 0.87 | 0.89 | 0.90 |
| Bad bond lengths & bad bond angles | 0/47361 (0%)<br>1/63882 (0%) | 0/14106<br>0/19056 | 0/20236<br>1/27315 | 0 / 61791<br>17 / 83443 |
| Molprobity score | 1.41 | 1.23 | 1.24 | 1.89 |
| Clashscore | 3.52 | 1.85 | 2.22 | 10.67 |
| Ramachandran plot Z-score | - 1.07 | -1.01 | -1 | 1.04 |
| Ramachandran Plot (%) | Outliers : 0.34 %<br>Allowed : 2.94 %<br>Favored : 96.73 % | Outliers : 0.18 %<br>Allowed : 3.99 %<br>Favored : 95.83 % | Outliers : 0.39 %<br>Allowed : 2.51 %<br>Favored : 97.09 % | Outliers: 0.04%<br>Favoured: 96.94% |
| Ramachandran rotamers (%) | Outliers : 1.20 %<br>Allowed : 5.51 %<br>Favored : 93.29 % | Outliers : 0.80 %<br>Allowed : 3.28 %<br>Favored : 95.92 % | Outliers : 1.26 %<br>Allowed : 4.93 %<br>Favored : 93.82 % | Outliers : 1.64 %<br>Favored : 76.22 % |
| CaBLAM outliers (%) | 1.5% | 1.1%o | 1.8% | 1.6% |
| EMRinger score | 2.86 (unsharpen)/3.19 (EMReady) | 3.07 (symmetrised) | 4.66 (symmetrised) | N/A |
